# The Impact of Genetic Background and Cell Lineage on the Level and Pattern of Position Effect Variegation

**DOI:** 10.1101/740951

**Authors:** Sidney H. Wang, Sarah C.R. Elgin

## Abstract

**Background:** Chromatin-based transcriptional silencing is often described as a stochastic process, largely because of the mosaic expression observed in position effect variegation (PEV), where a euchromatic reporter gene is juxtaposed with heterochromatin. Here we closely examine the impact of genetic background on PEV phenotypes in the fruit fly, *Drosophila melanogaster*.

**Results:** Using consecutive generations of selective breeding, we isolated, from a single laboratory population, two inbred lines exhibiting contrasting degrees of variegation (A1: low expression, D1: high expression). Within each inbred population, remarkable similarity is observed in both the degree and the pattern of variegation. 89.63% of the differences between the two inbred lines in the degree of silencing can be explained by genotype, while a modest but significant sex effect is also observed. Further analyses of the PEV phenotype in the progeny of crosses between A1 and D1 suggest that the genotypic effect is the result of the combined effect of multiple independent *trans-*acting loci. While the initial observations are based on a PEV phenotype scored in the fly eye (*hsp70-white* reporter), similar degrees of silencing were observed using a *beta-gal* reporter that can be scored across the whole fly. The pattern of variegating *hsp70-white* expression among individual flies becomes almost identical after five generations of inbreeding. Using a reporter inserted into the heterochromatic fourth chromosome, image analysis found significant enrichment of pigmentation in the ventral-posterior quadrant in both the A1 and D1 lines, and in the F1 and F2 progeny produced from a cross between A1 and D1, despite different degrees of expression.

**Conclusions:** Combining these results with the spreading model for local heterochromatin formation, we propose an augmented stochastic model to describe PEV. In this model, the genetic background, which determines the overall level of silencing, works with the cell lineage specific regulatory environment to determine the on/ off probability at the reporter locus in each cell. This model acknowledges cell-type specific events, as well as the general impact of heterochromatin formation.

## Background

Position effect variegation (PEV) describes the mosaic expression of a phenotype in a cell population that is otherwise thought to be uniform. It has generally been studied in cases where the cell-autonomous phenotype is easily visualized, such as eye pigmentation, but appears to be a general phenomenon [1,2]. Muller reported the original observation of variegating eye pigmentation in adult flies, recovered following X-ray mutagenesis. Because of the high degree of variation in the pattern, and in the level of pigmentation between individuals and across generations, he described the phenotype as “ever sporting” [3]. The report on this highly variable phenotype led to various speculative models describing how such a heritable, yet variable phenotype could arise [4]. Further investigations have led to a generally accepted transcriptional silencing model based on a stochastic spreading of heterochromatin [5]. The X-ray-induced inversion generated by Muller juxtaposed the *white* gene, which is required cell-autonomously for proper deposition of eye pigment, with the pericentric heterochromatin. The spreading of pericentric heterochromatin packaging to the *white* locus results in concomitant silencing; when this occurs in some but not all of the cells, the result is a variegated pattern of eye pigmentation. This spreading process is thought to be stochastic (reviewed in [6]).

Screens for second-site modifiers of a PEV phenotype have identified numerous loci that have a strong impact on the expression level of the PEV phenotype for the cell population as a whole. These genetic modifiers are referred to as suppressors [Su(var); loss of silencing]] or enhancers [E(var); gain in silencing] of PEV (see [2,6] for review). Some of these loci exhibit antipodal effects, i.e. if one dose of the gene results in loss of silencing, three doses results in an increase in silencing. This antipodal response has been interpreted as evidence that the probability of the heterochromatin spreading process occurring is determined at least in part by the dosage of key gene products that constitute the structural components of heterochromatin; in other cases an enzymatic contribution is implied. An assay for a PEV phenotype following a one generation cross to assess dominant effects of candidate PEV modifier alleles has therefore been commonly used to test the participation of a given gene of interest in the process of heterochromatin formation and gene silencing [7]. In fact, screens for PEV suppressors have been a major source of candidate genes for further analysis of the process of heterochromatin formation [8,9].

We previously devised a P element reporter to probe the heterochromatin landscape of the genome, P{*hsp26-pt, hsp70-white*}. Using the well-characterized *hsp70* promoter to drive a *white* reporter gene, about 1 % of the insertion lines recovered following mobilization exhibit a variegating eye phenotype [10]. Mapping of these variegating insertion lines revealed an outline of heterochromatin distribution in the genome, which is in agreement with prior cytological assignments, but provides higher resolution. PEV is observed following insertion of the reporter P element into the pericentric and telomeric regions of the major autosomes and the X chromosome, as well as regions of the Y chromosome. Based on the eye phenotype, the fourth chromosome (Muller F element), while largely heterochromatic, appears to have interspersed heterochromatic and permissive domains [11]. Characterization of these variegating P element reporter insertion lines has indicated that the basic principles for variegation as observed in the original *white* mottled line from Muller (i.e. sensitivity to sex chromosome dosage, etc.) are common to most variegating lines [10], although individual heterochromatic domains can show differences in sensitivity to a subset of the known suppressors of variegation [12–14]. A major exception are insertions into the TAS (Telomere Associated Sequences) sequences, just proximal to HeT-A and TART; these lines exhibit a PEV phenotype that is sensitive to mutations in the Polycomb silencing machinery [15], while ChIP analysis shows association with Pc [16]. Here we have used as our test locus a reporter in the fourth chromosome, a largely heterochromatic domain that for the most part mimics pericentric heterochromatin, a chromatin structure dependent on H3K9 methylation and HP1a [10,12].

Although PEV has been tremendously helpful in developing our understanding of heterochromatin, its stochastic nature continues to raise unanswered questions. Numerous mutations have been identified to modify PEV; it is estimated that there are more than 150 such modifiers in the fly genome [17]. Genetic background – including different assortment of alleles at these many loci – could affect the probability for a spreading event to occur in a given fly within a stock, and thus could contribute to the variation seen in PEV phenotypes. In fact, a recent study looking at PEV in an outbred fly population suggested that many more modifier loci likely exist across the genome [18]. Here, we examine the variation in PEV phenotype in a single laboratory reporter line using P{*hsp26-pt, hsp70-white*}. The study was carried out using a 4^th^ chromosome PEV reporter line, 39C-12, for several reasons. 39C-12 is relatively well-characterized in terms of its response to PEV modifiers [12,13,19]. Its position has been mapped to a precise location in the genome [11]. The *hsp70* promoter used in this reporter is well-characterized [20]. It’s basal activity at 25°C in this construct is sufficient to cause a uniform red eye when the reporter is inserted into a euchromatic site. Finally, the 39C-12 stock is considered relatively “clean,” because of the genetic bottleneck that occurred during the production of the transgenic line; specifically, the line is derived from a single male with the P element insertion on the fourth chromosome, back-crossed to *y w*^*67c23*^ females. The starting stock of 39C-12 had a high level of variation in PEV. We found that much of the variation in the strength of the PEV phenotype (i.e. the degree of silencing) for this stock could be attributed to genetic variants floating in the background. Intriguingly, we found fairly consistent patterns in PEV phenotype among individual flies across the inbred populations, indicating a more controlled expression of the PEV phenotype between cells than a purely stochastic heterochromatin spreading model would otherwise suggest. Our observations resonate well with published literature and provide fresh insights into this classic system.

## Results

Visual inspection shows considerable variation in the levels of PEV in adult fly eyes among individuals of the 39C-12 reporter line raised at 25°C. Despite a genetic bottleneck during the production of this transgenic line, there is a high degree of variation in the level of extracted eye pigment (coefficient of variation [CV] = 51.3%), which averaged 0.0246 (OD_480_). To study the underlying genetic contribution to the variation of PEV among individual flies, we selected for extreme PEV phenotypes (by eye pigment levels). A single virgin female from the parental population displaying the strongest PEV eye phenotype was mated to a single male sibling with a matching eye phenotype. This process was repeated for 5 generations (i.e. full sibling mating followed by selection) to obtain a fly line, 39C-12-A1, in which the level of eye pigmentation is lower (i.e. strong PEV; mean = 0.0104; SD = 0.0020) and more consistent among individuals (CV = 19.23%; Figure 1; Table 1) than the starting population. A weak PEV line, 39C-12-D1, was similarly derived (Figure 1; mean= 0.0345; SD= 0.0076; CV = 22.03%). The two lines have 3.3-fold pigment level difference (p < 1e-11, ANOVA) and represent the extreme ends of the phenotypic spectrum of the original population (i.e., each is about one standard deviation away from the mean of the original 39C-12 population, in opposite directions). In addition to the genetic effect introduced by selective breeding, there is also a sex effect impacting the PEV phenotype. While there is a modest 26.02% higher pigment level in A1 males (relative to A1 females, p < 0.05), there is a 47.93% higher pigment level in D1 males compare to D1 females (p < 0.001). Overall, the combined genotype and sex effects explain 95.9% of the variance between A1 and D1 flies, while genotype alone explains 89.6% of the variance.

**Table 1.**
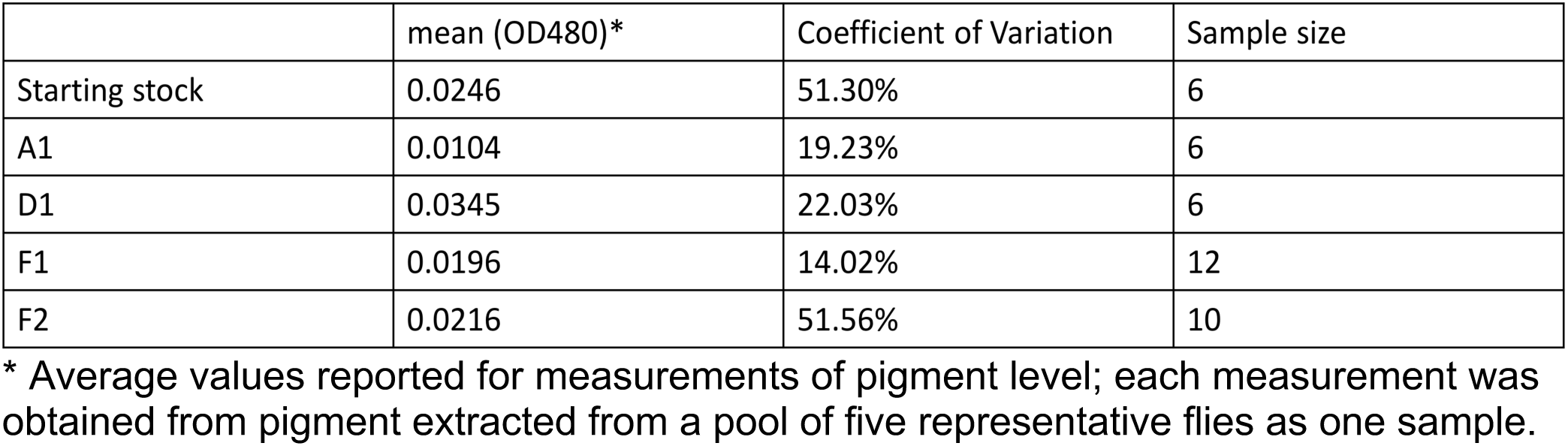
Pigment assay results for 39C-12 inbred variegating lines

**Figure 1.**
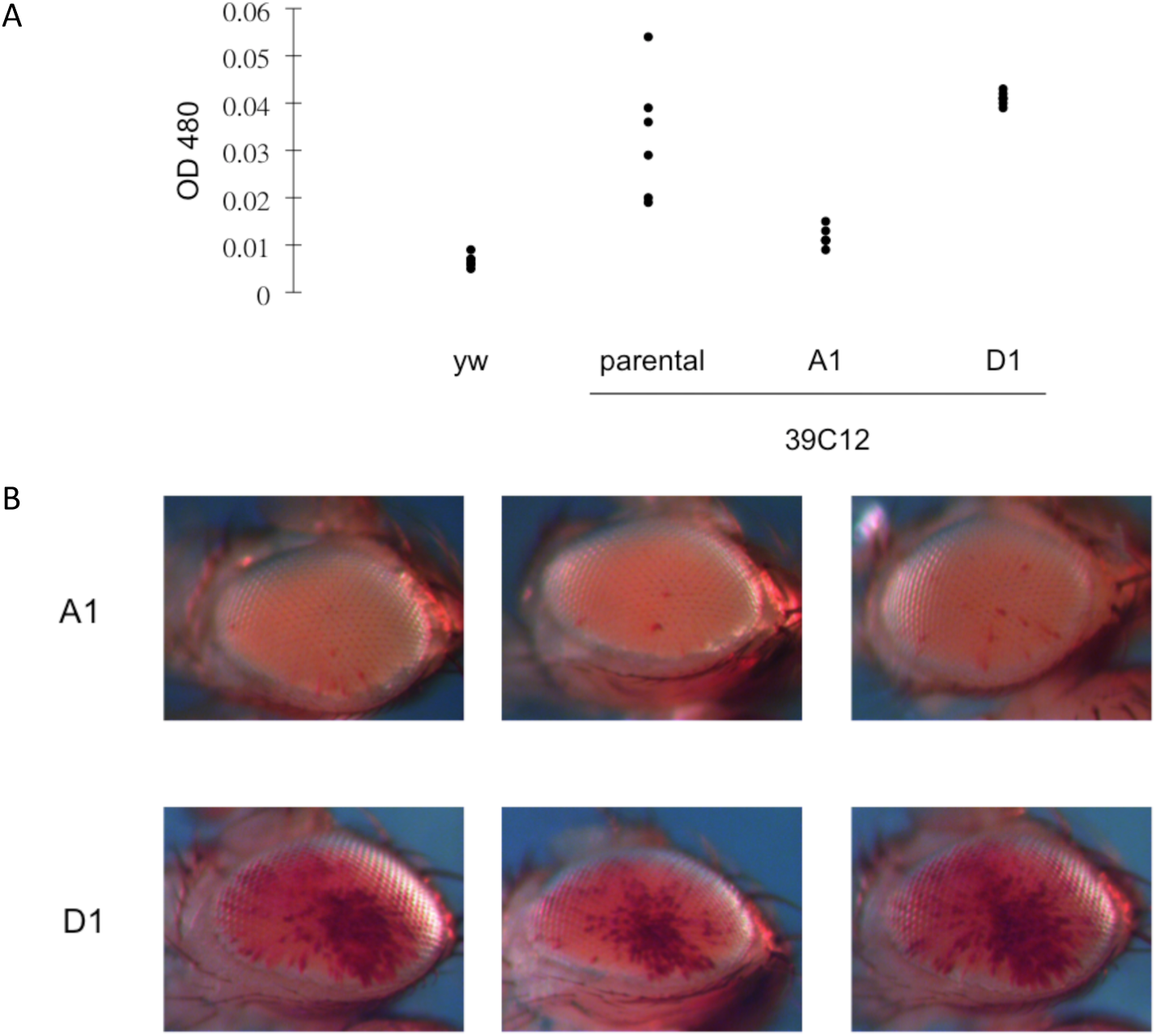
Selective inbreeding results in highly consistent PEV phenotypes within a laboratory population. (A) Quantitative assessment of pigment levels in the adult fly eye representing the degree of PEV. Each data point represents a reading from samples of five flies from a population of the indicated genotype, parental (39C-12) or selected (A1, D1) (see Materials and Methods for details). *yw* is used to indicate the background pigment level. (B) Images of the PEV pattern in the adult fly eye taken from randomly selected individuals in each of the A1 and D1 inbred populations.

To assess the genetic architecture of the two inbred lines regarding its impact on the PEV phenotype, crosses were performed between these lines to generate F1 and F2 populations. There is a fairly consistent intermediate PEV phenotype in the F1 population (mean = 0.0196, SD = 0.0027, CV = 14.02%). Similar results were obtained from crosses in both directions (Fig 2 A, B). The average pigmentation level for F1 progeny in both cases falls right in the middle between the pigment levels of the parental A1 and D1 lines (Fig 1A, 2A; Table 1). As would be expected for a quantitative trait involving multiple independent loci, a wide spectrum of PEV phenotypes was observed in the F2 population (mean = 0.0216, SD = 0.0111, CV = 51.56%), which likely resulted from meiotic recombination and random segregation of the A1 and D1 background PEV modifiers. The mean pigmentation level for the F2 progeny is similar to the F1 population (0.0216 vs. 0.0196); in contrast, there is a large increase in the range of expression levels for the PEV phenotype between individuals of the F2 population (CV = 51.56% vs. 14.02%; Fig 2A, 2C). The range of phenotypic variation in the F2 population resembles that of the starting 39C-12 stock (compare Fig 1A with Fig 2A, CV = 51.3% vs. 51.56%). Taken together, these results suggest that the variation in PEV phenotype between individuals of the 39C-12 transgenic fly line is best described by the effect of multiple *trans* genetic modifier loci acting independently in the background.

**Figure 2.**
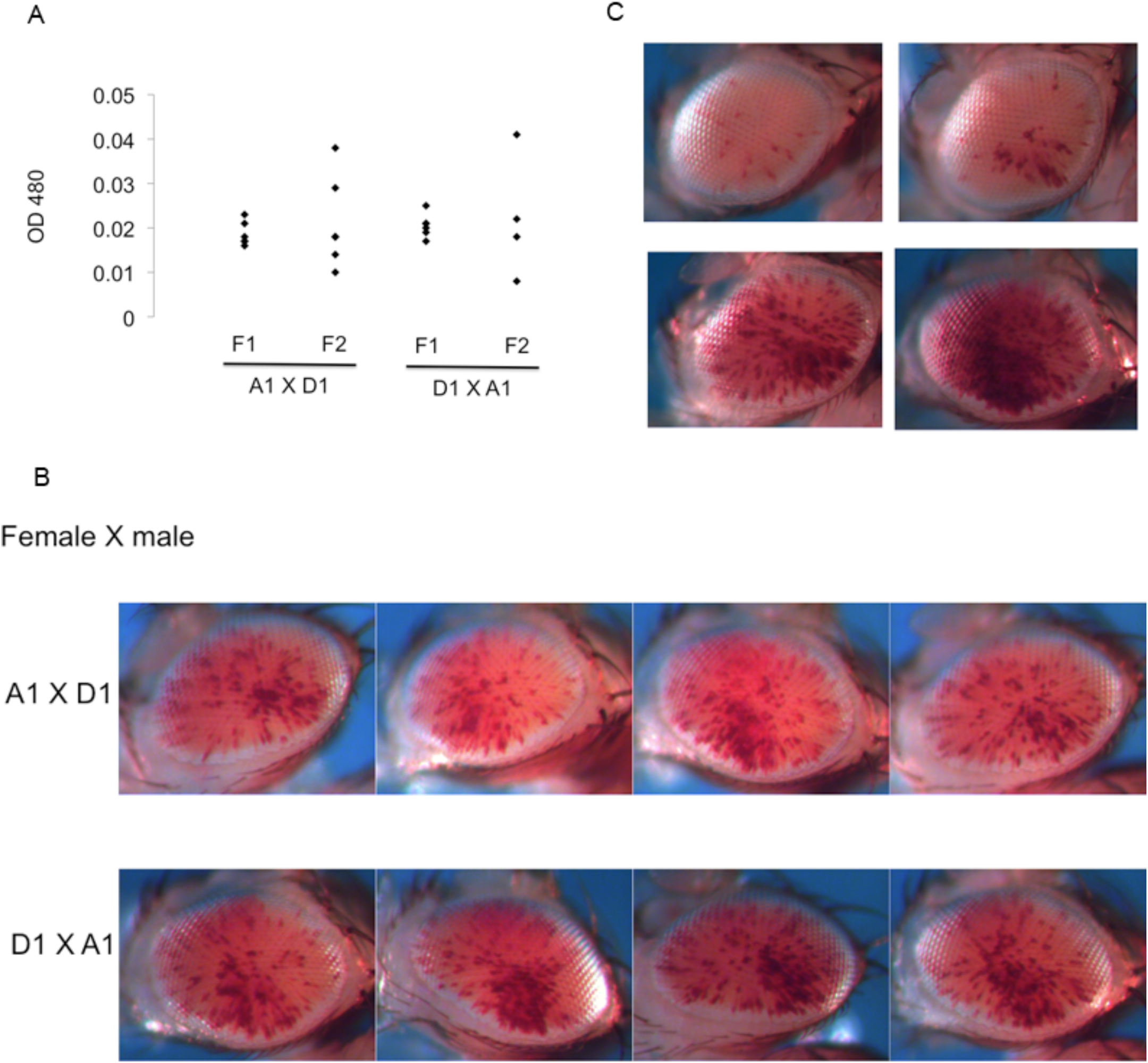
PEV phenotype of the progeny from the cross between the A1 and D1 inbred lines. (A) PEV levels in the adult progeny. Each data point represents a sample of five flies from a population of the indicated genotype (see Materials and Methods). Results observed were essentially the same from crosses in either direction (females listed first). (B) The PEV pattern in the adult fly eye from randomly selected F1 progeny of a cross between the A1 and D1 inbred lines in the indicated direction. (C) Selected images of the PEV pattern in the F2 population representative of the diversity in pigmentation levels observed.

The results above are based on the PEV eye phenotype of a P-element *hsp70*-*white* reporter inserted into the heterochromatic 4^th^ chromosome. To determine whether the conclusions drawn are generally applicable to the PEV phenotype, we evaluated the impact of the A1 and D1 background genotypes on the PEV phenotype of a Y-linked *hsp70-LacZ* PEV reporter, *Tp(3;Y)BL2* (BL2). The PEV phenotype of the BL2 *LacZ* reporter line used for this purpose results from a translocation of a fragment of the 3^rd^ chromosome carrying the reporter to the Y chromosome following X-ray irradiation [21]. A multigeneration cross scheme was designed to introduce this Y-linked PEV reporter into the A1 (or D1) genotype, without perturbing that genetic background, utilizing dominantly marked balancer chromosomes (and the fact that meiotic recombination is not known to occur in the male germ line [22,23]) (Supp. Fig 1). Two inbred lines containing the Y-linked BL2 reporter in the A1 and D1 genetic backgrounds, respectively, were derived (BL2-A1 and BL2-D1). The level of *beta*-galactosidase activity (mAU/min) in lysates prepared from single male flies was used as a quantitative readout for the PEV phenotype. A consistent PEV phenotype for the BL2 reporter across individuals from each of the A1 or D1 genetic backgrounds was observed, with D1 flies exhibiting ∼4.64 times the activity of A1 flies (Fig 3). The BL2 reporter in a D1 background gave a CV of 13.11% (mean = 1.44, SD = 0.19), while in an A1 background it gave a CV of 17.24% (mean = 0.31, SD = 0.05). Taken together, the results for the BL2 reporter largely recapitulate the results for the 39C-12 reporter, indicating that the same (or similar) background PEV modifiers impact both a 4^th^ chromosome P-element PEV reporter and an X-ray induced Y-chromosome-linked PEV reporter in the same direction. Lu et al. reported variegating expression for the BL2 reporter in multiple tissues, such as various differentiating imaginal discs in late third instar larvae, as well as in adult eyes [24]. Here, BL2 PEV was surveyed using the whole fly in a quantitative assay. Given the Lu et al. results, the findings here generalize the impact of background modifiers on PEV (i.e., heterochromatic silencing) beyond the fly eyes analyzed using 39C-12. It is noteworthy that insertion of the same P element reporter into different heterochromatic domains results in different PEV phenotypes, with the “salt and pepper” pattern commonly associated with insertions into pericentric heterochromatin and the fourth chromosome, while large patch or sectored mosaicism is associated with the Y chromosome [10,21]. The results above demonstrate similar quantitative responses to PEV modifiers regardless of the over-all geometry of the expression pattern.

**Figure 3.**
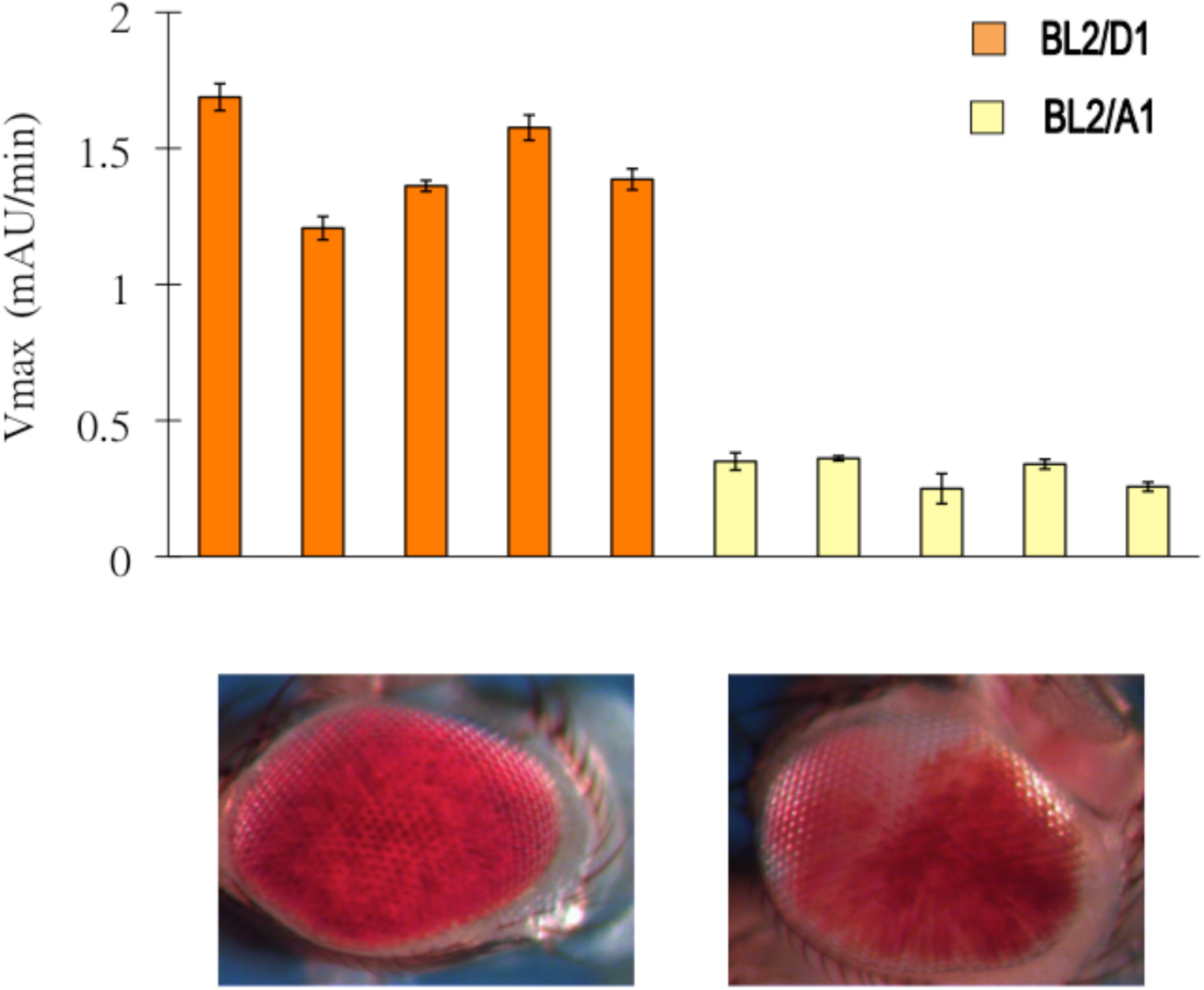
PEV phenotype of the Y-linked BL2 reporter in the A1 and D1 genetic backgrounds. The level of PEV is quantified by measuring the activity of the *beta*-galactosidase reporter gene. Each bar represents the activity level measured in lysate prepared from one adult whole fly of the indicated genotype. Bar height and error range represents the mean and standard error calculated from technical replications (i.e. measurements made on aliquots of the same lysate). Representative images of eye pigmentation for each genotype, shown below the bar graph, show the variegating phenotype anticipated.

The results on the strength of PEV shown above suggest a strong genetic component in determination of the overall level of heterochromatic silencing, which could be explained by background genetic variants impacting PEV modifiers. In addition to a consistent level of eye pigmentation, the inbred lines also showed a more consistent pattern of eye pigmentation among individuals within each line (Figure 1B). An enrichment of pigmented ommatidia in a ventral-posterior sector (i.e. near the neck) is observed for both A1 and D1 lines. The pigment enrichment pattern was evaluated by comparing the proportion of ommatidia exhibiting red pigment between the ventral-posterior quadrant of the eye and the rest of the eye (Supp. Fig 2). For the A1 and D1 lines, the ommatidia in the ventral-posterior quadrant are 5.24 times (p < 0.05, ANOVA, n = 5) and 3.58 times (p < 1e-5, ANOVA, n = 5) more likely to exhibit pigment than ommatidia in the rest of the eye. Furthermore, in both F1 and F2 progeny of an A1 by D1 cross, where no manual selection for eye phenotype was done, significant enrichment of the pigment level is observed in the ventral-posterior quadrant of the fly eye. F1 progeny exhibit a 2.85-fold enrichment of pigmentation in the ventral-posterior quadrant (p < 1e-6, ANOVA, n = 14), while F2 progeny exhibit a 2.72-fold enrichment of pigmentation in the ventral-posterior quadrant (p < 0.05, ANOVA, n = 10). This consistent enrichment (see Table 2 for a summary) suggests differences in the ability of cells in the ventral-posterior quadrant and cells in the rest of the eye to silence the reporter. These results argue that nonrandom variation in heterochromatin silencing among cells plays a role in determining the observed pattern.

**Table 2.** Image analysis results for 39C-12 inbred variegating lines

|  | Fold enrichment (i.e. in/out)* | Significance | Number of images analyzed |
| --- | --- | --- | --- |
| A1 | 5.24 | $p < 0.05$ | 5 |
| D1 | 3.58 | $p < 1e-5$ | 5 |
| F1 | 2.85 | $p < 1e-6$ | 14 |
| F2 | 2.72 | $p < 0.05$ | 10 |

\*Fold enrichment indicates the proportion of pigmented pixels within the ventral-posterior quadrant vs. the proportion of pigmented pixels outside that quadrant.

## Discussion

A high level of heterogeneity in PEV phenotypes is commonly observed within laboratory reporter strains, even when the genetic background is standardized (as was the case here) by back crossing to *yw*^*67c23*^. Through selective breeding, we successfully derived inbred fly lines from the 39C-12 parent stock that show consistently strong or weak PEV (lines A1 and D1). Further analysis of the PEV phenotype from a two-generation cross between the two inbred lines indicates that there are likely multiple *trans* genetic modifier loci in the background acting independently to determine the strength of PEV. This conclusion is reinforced by results from analyzing the Y-linked BL2 reporter; there, a similar impact of the A1 or D1 genotype was observed on a different reporter, which removes the possibility of *cis*-linkage between the reporter and the background PEV modifier variants. These results resonate well with a recent study on PEV phenotype using an outbred population [18]. There, Kelsey and Clark found a large number of genetic variants across the genome significantly associated with the strength of PEV. Here, we have demonstrated that even within a relatively inbred population, one still observes a high level of variation in phenotype, which is best described by the effects from multiple independent modifier loci.

While this study mostly focused on the genetic background of a single reporter line, 39C-12, this level of heterogeneity in PEV phenotype is commonly observed in our laboratory across multiple reporter lines. Given that we successfully derived two inbred lines that have more than 3-fold difference in eye pigment level from a single parent population, it is worthwhile considering the implications for using a reverse genetic candidate approach for identifying novel PEV modifiers [2,6]. In addition to forward genetic screens, novel PEV modifiers are often identified through a reverse genetic screen for dominant effects on PEV using mutant alleles of genes that are suspected to play a role in chromatin-based transcriptional silencing (e.g. SETDB1, G9a, PIWI). High levels of variation in the starting reporter PEV line often result in high levels of variation in the readout for the screen; some researchers will therefore decide to homogenize the genetic background of the starting reporter line to reduce variation in order to gain power to detect dominant effects from the mutant alleles being tested. In light of the results presented here, the steps researchers take to homogenize the background could lead them to unwittingly enrich or deplete the genetic background of variants that might interact with the mutant allele of interest, and therefore amplify or reduce the phenotypic impact. This practice could therefore lead to inconsistent findings between labs and potentially misleading results. For more reproducible findings, instead of homogenizing the genetic background, researchers could increase power by increasing sample size. This suggestion is particularly appropriate for model organism researchers, such as fly geneticists, considering the limited cost imposed by increasing sample size for a pigment assay or an enzymatic assay for assessing PEV phenotype, even by 5 or 10 times the current standard.

The results using the BL2 reporter on the Y chromosome reproduced the results observed using the original 4^th^ chromosome P element insertion reporter, in that the Y chromosome reporter showed the same direction of response at similar magnitude to the collection of variants in the A1 and D1 genetic background. Assuming that these results from one Y chromosome reporter and one 4^th^ chromosome reporter are representative of Y chromosome and 4^th^ chromosome heterochromatin (i.e. given the caveat of small sample size and the limitation that both reporters use a 5’ regulatory region of an *hsp70* gene), the results indicate that the background modifiers in aggregate have similar effects on the PEV phenotype, and by extension, on heterochromatic silencing, of two different chromosomes. This conclusion resonates with many previous studies indicating shared PEV modifiers (see [6] for review).

When considered in conjunction with the mass action model (proposed by Locke et al. and widely accepted) [25], the results are consistent with a model describing the Y chromosome as a heterochromatic sink that modifies PEV phenotype of reporters at other genomic loci by trapping structural protein products of PEV modifiers. This effect of Y chromosomes, in light of the mass action model, potentially explains the sex-linked impact on PEV observed here and previously reported in many investigations.

Moreover, the BL2 reporter made it possible to survey the PEV phenotype from the entire fly, which further generalizes these conclusions. However, in contrast to the sharing of modifier effects observed here (and elsewhere) between the Y and the 4^th^ heterochromatin, it is amply documented that different Su(var) mutations can have a different impact on different heterochromatic domains (e.g. Brower-Toland et al, 2009; Phalke et al 2009; Wang et al 2014) [13,14,26]. Thus, a future study extending to evaluate the impact of the same (or similar) genetic background on multiple reporters inserted in different heterochromatic domains across the genome is likely to reveal new insights on the extent of sharing across domains.

A rather consistent pattern of eye phenotype became better defined as the flies became increasingly inbred over generations of full sibling crosses. Using the shape of the fly eye and other anatomical structures surrounding the eye as landmarks, enrichment of pigmentation at the ventral-posterior quadrant of the eye was tested. We found significant enrichment for A1 and D1 lines, as well as the F1 and F2 progeny of an A1 by D1 cross. It is not uncommon among fly PEV researchers to observe certain patterns consistently occurring in certain reporter lines. For example, insertions of the *hsp70-white* reporter into the Y chromosome often show patterns with large blotches of pigmentation, whereas insertions of the same reporter into pericentric heterochromatin or the fourth chromosome often results in a fine-scale salt-and-pepper appearance in the eye [10,26]. This study followed up on observations of a PEV pattern in the 39C-12 reporter line through selective breeding and then formally tested the enrichment pattern through statistical analysis on images of the PEV phenotype. The results suggest a more controlled reporter expression than would be inferred from a random spreading of heterochromatin. While the random spreading model is adequate for explaining sectors of PEV reporter expression in *S. pombe* colonies [1,5], in higher eukaryotes, it appears that a more elaborate model considering developmental lineage differences between cells displaying the phenotype is needed to adequately describe the process.

Consistent with this conjecture, we did not find the Y chromosome BL2 reporter, which is in a different cis-regulatory environment and likely subject to different regulation during the process of developmental lineage differentiation, to express similar patterns of PEV phenotype as the 39C-12 reporter, despite having been transferred into the same A1 or D1 genetic background. One suggestion as to the source of differences in PEV patterns between cell types is the timing of the last wave of cell divisions during metamorphosis. Using the beta-gal reporter, Lu et al. reported that in the eye disk, there is a dramatic difference in variegation on either side of the morphogenetic furrow, with a relaxation of silencing at this juncture [24]. Such features could be a contributor, as cells that go through an S phase later in developmental time are inherently exposed to a different environment during this process.

## Conclusions

In summary, our observations with two PEV reporters inserted in different *cis*-regulatory environments in two diverging *trans*-acting genetic backgrounds indicate that while *trans*-acting modifiers play the major role in determining the degree of PEV silencing, the pattern of PEV silencing is likely influenced more by the *cis*-regulatory environment of the reporter insertion site.

## Materials and Methods

### Fly husbandry and genetics

Flies were cultured at 25°C, 70% humidity on regular cornmeal sucrose-based medium [27]. Unless otherwise specified, genetic crosses were performed by mating 2 male flies to 3∼5 female virgin flies. The 39C-12 reporter line [10] was used as the starting line to generate A1 and D1 inbred lines. Five generations of consecutive full sibling crosses with selection for extreme eye phenotype at each generation were performed to create the two inbred lines. To substitute the BL2 Y-linked PEV reporter [21] into the A1 or D1 genetic background, dominantly marked balancers were used to follow the second and third chromosomes (see Supp. Fig 1 for crossing scheme). Balancers SM5 and TM6 were first introduced to the BL2 line by a standard cross, and the F1 male progeny that had both second and third chromosomes balanced were selected (based on the dominant markers) to mate with female flies from the inbred line. A male F2 progeny from the F1 cross with both balancers over inbred chromosomes were selected to backcross to 3∼5 inbred line female virgins. Only one male fly was used in the F2 cross in order to ensure that there would only be two genotypes of 4^th^ chromosome in the F3 population (i.e. the original 39C-12 4^th^ chromosome and the other 4^th^ chromosome, which is denoted as +^iso^ in Supp. Fig 1, carried by the selected F2 male). Because the 4^th^ chromosome is not known to recombine during meiosis (or does so extremely rarely), progeny from the backcross lacking both balancers were selected to make a floating stock (i.e. inbred genetic background with an unmarked 4^th^ chromosome floating). This floating stock was made homozygous for the 39C-12 4^th^ chromosome, as judged by the presence of the 39C-12 reporter expression, to generate the final stock. In order to evaluate the homozygosity of the 4^th^ chromosome in the final stock, 39C-12 reporter expression in all female progeny was followed by visual inspection for multiple generations. (*white* expression in male progeny cannot be used to evaluate 39C-12 expression because of the interference coming from the mini-*white* construct in the BL2 reporter.)

### PEV assays

Eye pigment extraction and quantification was done essentially as previously described [28] with a few modifications. Instead of hand homogenizing for pigment extraction, whole flies were homogenized using a Mixer Mill Mm 300 to increase the throughput and consistency. The overnight incubation at 4°C was omitted. For each genotype of each sex, 20∼30 age-matched flies (3∼5-day-old) were randomly selected from the population and sorted according to their pigmentation level by visual inspection. Five flies of similar pigmentation levels were than collected together as one sample. The same protocol was used for both sex.

X-gal staining of eye imaginal discs and the assay of *beta*–galactosidase activity were carried out as previously described [29].

For image analysis of the PEV phenotype, eye pictures from a random sampling of the PEV phenotype from each population were selected based on eye size and angle of the photo to ensure consistent quantification. Each image was then converted to 8-bit grey scale in imageJ, and then further converted to a binary image through manual threshold setting. The guideline used for threshold setting was to capture as many variegating speckles as possible without introducing large patches of shadow that resulted from lighting differences during imaging. The area to quantify was manually selected using the imageJ *oval tool* to cover as much of the eye as possible without covering other anatomical structures. The binary oval image representation of the eye PEV phenotype was then converted into a binary table of quantification using the imageJ *image to result* function, where each entry in the table represents a pixel in the image. To test for enrichment of pigmentation in the ventral-posterior quadrant, the proportion of “expressed” pixels in the ventral-posterior quadrant was compared to the proportion of “expressed” pixels outside of the quadrant of interest using statistical tests described in the following section.

### Statistical analysis

Analysis of the strength of the PEV phenotype (e.g. pigment level) was done using either R or excel. For estimating the genotype effect and the sex effect on the variation in PEV phenotype between the A1 and D1 lines, we fit the pigment assay data to a linear model using the genotype label and sex label as predictors. More precisely, the OD 480 reading for pigment level was first log transformed and then fitted as the response variable in the following linear model using the lm() function in R:

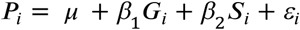

where *P*_*i*_ represents the pigment level of individual *i, G*_*i*_ is the indicator variable for the genotype label, and *S*_*i*_ is the indicator variable for the sex label. The adjusted R^2 produced by applying the summary() function to the above described lm object is used to estimate the percentage variance explained by the model. To evaluate the significance of the genotype effect (i.e. the differences in PEV phenotype between the A1 and D1 genotypes) we performed an F test by applying the anova() function on the lm object. Analysis of the pattern of PEV followed a similar linear model framework, where the proportion of pigmentation in or out of the ventral-posterior quadrant is modeled using a binary predictor of location. To evaluate the association between the proportion of pigmentation and the location we performed an F test.

## Declarations

### Ethics approval and consent to participate

Not applicable

### Consent for publication

Not applicable

### Availability of data and material

The data used and/or analyzed during the current study are available from the corresponding author on reasonable request.

### Authors’ contributions

SHW conceived the idea for this study. SHW and SCRE designed the study. SHW executed the experiments and data collection. SHW analyzed the data. SHW and SCRE interpreted the results and wrote the manuscript.

## Acknowledgements

We thank members of the Elgin lab, Joel Eissenberg, and Ian Duncan for their critical review of the manuscript. This work has been supported by a Howard A. Schneiderman Fellowship (SHW) and by NIH grant GM068388 (to SCRE).

## Supplemental Figure legends

**Supp. Figure 1.**
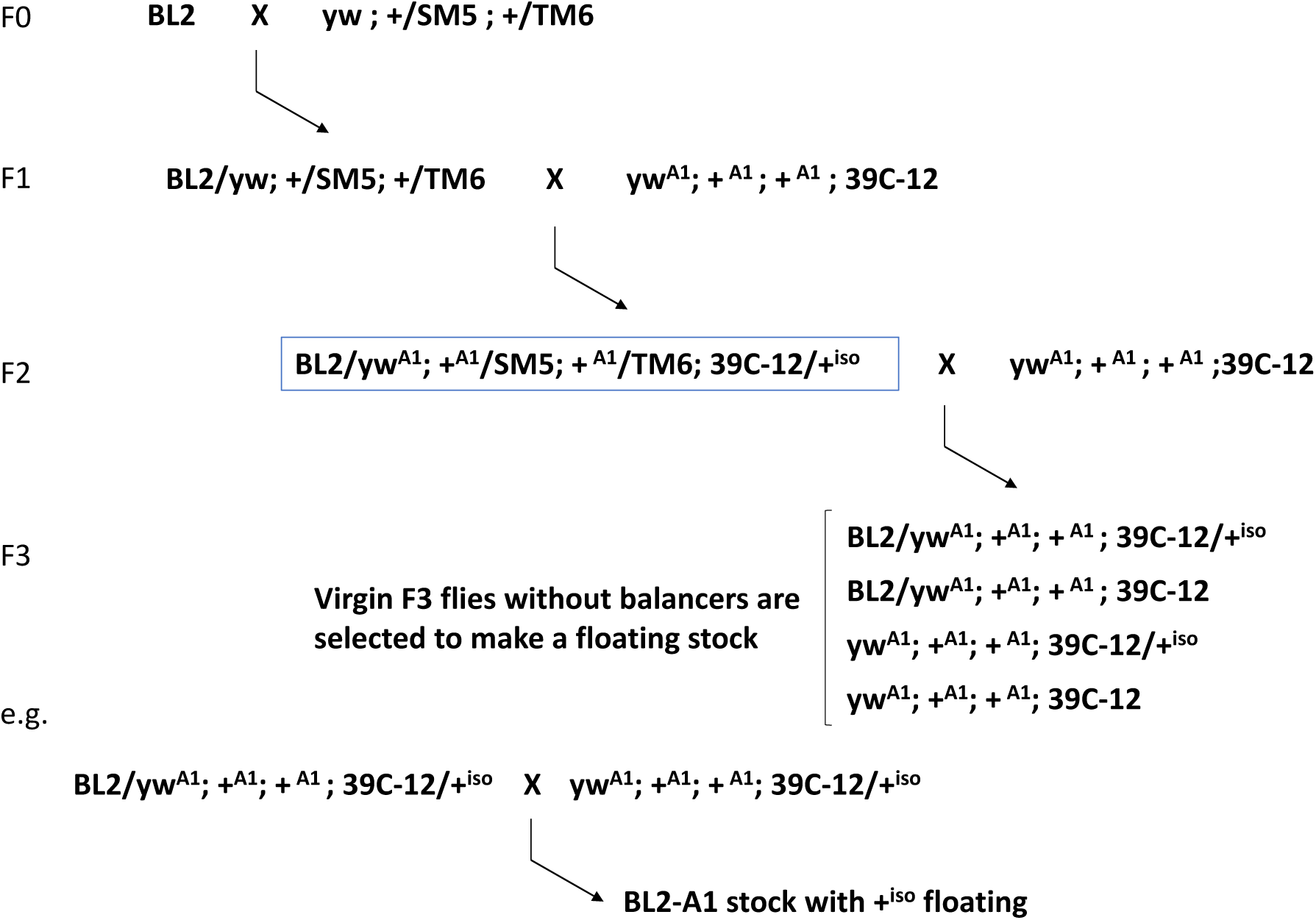
Crossing scheme for creating isogenic BL2 reporter lines. The BL2 reporter on the Y chromosome was first crossed into a balancer stock to recover second and the third chromosome dominant markers. The F1 male progeny with second and third chromosomes dominantly marked were selected to cross with female virgins of the A1 inbred line. A single F2 male progeny (blue rectangle) was selected to back cross to 3∼5 A1 inbred line female virgins. The F3 progeny that have no balancer chromosomes will have the BL2 reporter in the A1 background with a non-A1 4^th^ chromosome floating in the population. Note that the non-A1 4^th^ chromosome was introduced from a single F2 male, which means that in the F3 population there are only two genotypes of the 4^th^ chromosome (denoted 39C-12 and +^iso^ respectively). The F3 virgin flies that have no balancer chromosomes were selected to create a floating stock. In order to separate the 39C-12 chromosome from +^iso^ chromosome, single sibling pairs from the F3 floating stock were isolated to create multiple stocks; the 39C-12 reporter expression in all female fly eyes was followed by visual inspection for several generations in order to identify a homozygous 39C-12 stock (i.e. the exact A1 background with the Y chromosome containing the BL2 reporter). The same approach was used to transfer the BL2 reporter to the D1 genetic background.

**Supp. Figure 2.**
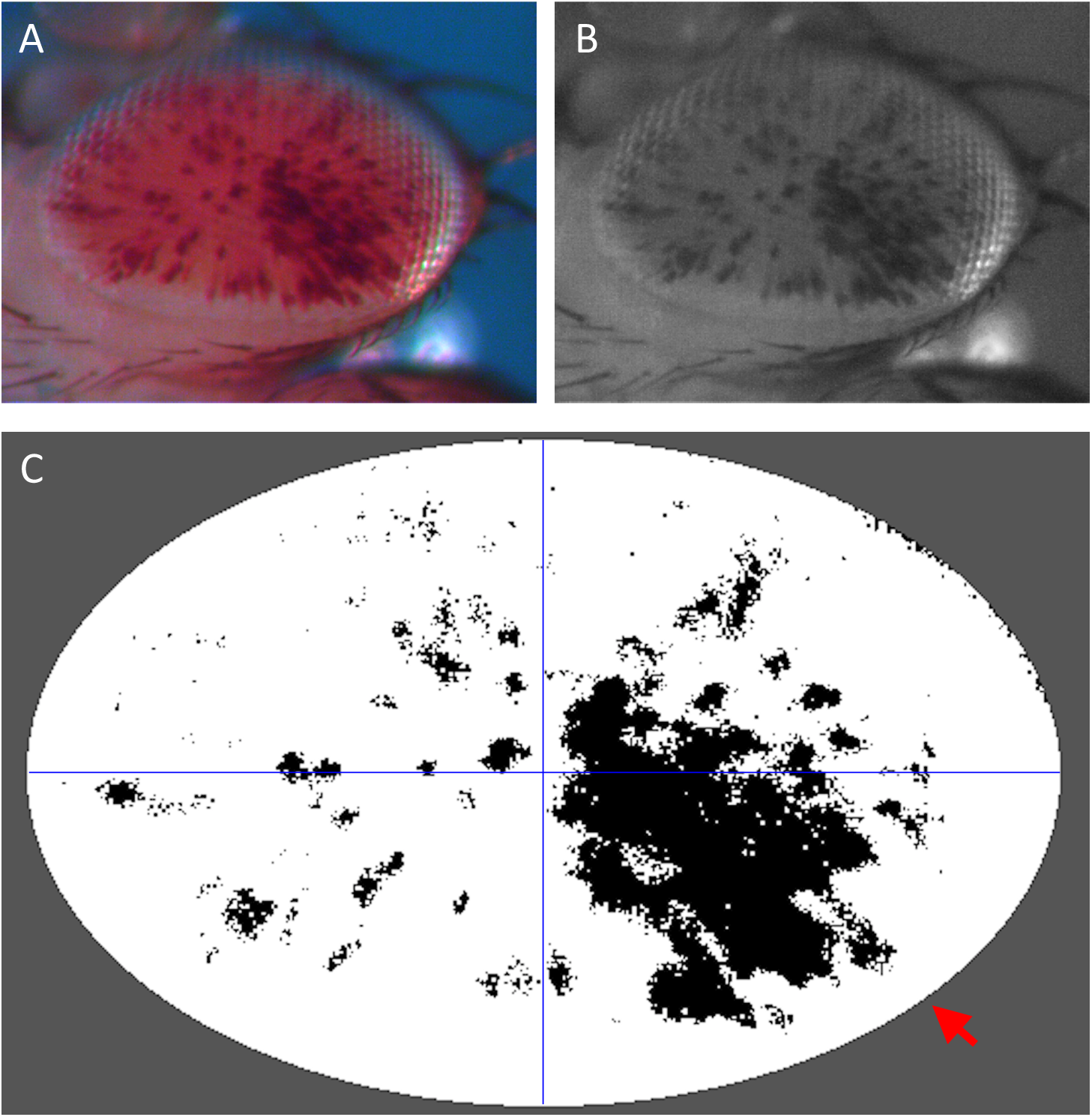
Example Images to illustrate the processing steps for quantifying the pattern of PEV. (A) The original photo of a representative variegating eye phenotype taken from an F1 male progeny of an A1 by D1 cross. (B) An 8-bit grey scale version of A transformed using imageJ. (C) A binary image of B generated using imageJ. The image was first rotated so that the maximal area of the fly eye could be selected using the oval tool. Pixels outside the selected oval area were removed (pseudo-colored in grey for illustration) while pixels within the oval area were converted to binary by setting a threshold. The threshold selection was done manually to best represent the original eye phenotype. To evaluate the similarity of the PEV pattern between individuals, each image of a fly eye was split into four even quadrants (blue lines) and the pigment enrichment (i.e. the proportion of black pixels) in the ventral-posterior quadrant (red arrow) was evaluated against the area outside the ventral-posterior quadrant.

